# SurfDiff: protein surface profiling for selective or broadly reactive epitope prioritisation in binder and immunogen design

**DOI:** 10.64898/2025.12.23.696170

**Authors:** Benedikt Jedlitzke, Oliver Wissett, Tung H. Nguyen, Elodie Ghedin, Pietro Sormanni

## Abstract

Rational selection of protein-surface patches – epitopes or ligandable sites – is essential for developing targeted antibodies, binding proteins, peptides, or small-molecule ligands, and for informing immunogen design in vaccines. We present SurfDiff, a structure- and sequence-informed framework for protein surface comparison. SurfDiff supports one-to-one and one-to-many comparisons by combining local structural alignments with physicochemically and spatially aware neighbourhood analysis. It assigns residue-level uniqueness and similarity scores, which can be aggregated into surface-level similarity, selectivity, and a discriminability score that captures conservation across desired targets while penalising similarity to undesired off-targets. These scores, calculated solely from the targets without any knowledge of the binders, reliably predict known experimental selectivity for diverse binders including small molecules, peptides, antibodies and antibody mimetics across viral antigens, cytokine isoforms, GPCR subtypes and serum albumin. They also correlate strongly with binding and neutralisation data across pathogen variants, and substantially outperform commonly used bioinformatic metrics such as sequence substitution matrices and structural similarity measures. SurfDiff provides a generalisable and interpretable approach to protein surface profiling, enabling selective or cross-reactive binder design, cross-species prioritisation, and facilitating immunogen selection. We make SurfDiff available as downloadable open-source software: (https://gitlab.developers.cam.ac.uk/ch/sormanni/surfdiff) and as a webserver: www-cohsoftware.ch.cam.ac.uk/index.php/surfdiff.

## Introduction

Selective biomolecular recognition is governed by precisely defined surface features. Identifying which regions of a protein surface can drive selective versus cross-reactive binding is central to the design of binders^1^ – here encompassing antibodies, antibody-mimetics such as mini-proteins, peptides, small-molecule ligands, etc. – and equally important for immunogen design^2^, where conserved, protective epitopes must be prioritised while avoiding off-targets and host liabilities. Robust epitope choice underpins downstream decisions on antigen engineering^2^, adjuvantation^3^, dosing and delivery^4^, and determines whether binding is narrow and specific^5,6^ or intentionally broad across variants^7^, isoforms^8^ or species^9^.

Recent advances in computational structural biology are transforming drug discovery and vaccine development^10–13^. Historically, delineating binding sites required structural determination^14^ or complex experimental screenings, including targeted mutagenesis^15^, hydrogen-deuterium exchange mass spectrometry^16^, and extensive competition and cross-reactivity assays^17^ – efforts often at least as labour-intensive as broad functional screening of hits. Conversely, it is now increasingly feasible to design binders tailored to predetermined epitopes of interest^12,13,18–23^ and to reliably predict binder-target complex structures^11,24,25^, enabling high-throughput and precise epitope mapping from the earliest stages of a programme. Yet, despite these developments, the fundamental question of which epitope(s) on a target protein are most suitable for the intended selectivity profile remains substantially under-investigated.

The answer can be straightforward for well-characterised targets with validated functional sites, but becomes non-trivial when there is a need to discriminate within families of closely related proteins^26^, or to balance binding breadth against avoidance of undesired off-targets^27,28^ or host proteins^29^. Similarly, in rapidly evolving pathogens that can evade previous immunity, such as SARS-CoV-2, the challenge is to prioritise developing therapeutics for broad neutralization by focussing on conserved pathogen epitopes.^30^

These scenarios highlight the need for a rational framework for epitope or binding site selection. This framework should: (i) operate at single-residue resolution to yield precisely defined candidate binding sites; (ii) support one-to-one and one-to-many comparisons across proteins – including proteins that are structurally dissimilar at the global level; and (iii) enable the quantification of distinct design objectives – identifying unique epitopes for selective binding, shared epitopes for broad binding, and intermediate cases where engagement of some targets but avoidance of others is desired. Furthermore, such framework should be interpretable and must rely solely on the knowledge of desired target(s) and undesired off-targets, independent of binder structure or affinity, thus enabling the pre-emptive evaluation of targets or antigens before committing to specific binder modalities or affinity/kinetic profiles.

To address these needs, we introduce SurfDiff, a structure- and sequence-informed method for systematic protein surface comparison. SurfDiff combines local structural alignment with physicochemically and spatially aware neighbourhood analysis to assign residue-level uniqueness and similarity scores. These are aggregated into surface-level similarity, selectivity, and a discriminability score that captures conservation across desired targets while penalising similarity to undesired off-targets. By providing quantitative, interpretable surface profiles independent of binder structure or affinity, SurfDiff is intended to guide epitope prioritisation for selective or cross-reactive binder design, cross-species applications, and immunogen selection.

In drug discovery, research and development programmes typically define a target product profile (TPP) and a candidate drug target profile (CDTP) that sets acceptance criteria for functionality, specificity, species cross-reactivity, binding affinity, and developability^31^. SurfDiff supplies quantitative, residue-level comparative metrics (similarity, selectivity and discriminability) that can streamline and sharpen the definition of TPP and CDTP criteria. More decisively, it makes possible a priori, pre-emptive, identification and ranking of target epitopes or binding sites most likely to deliver the desired profile, before committing to modalities or large-scale screening campaigns. These binder-agnostic metrics can be mapped directly to CDTP criteria – for example, prioritising epitopes expected to maximise on-target engagement while avoiding closely related off-targets, maximising the chances of achieving the species cross-reactivity required for pharmacology and safety studies, and maintaining coverage across human haplotypes where sequence polymorphism is relevant. By furnishing an interpretable, auditable rationale for site selection, SurfDiff shifts decision-making upstream, thereby reducing reliance on extensive, iterative laboratory screening that can be time-consuming, resource-intensive, and sometimes inconclusive.

## Results

### The SurfDiff pipeline for protein surface comparison

The SurfDiff pipeline enables single-residue-resolution comparisons of protein surfaces by integrating local structural alignment with neighbourhood-based physicochemical environment analysis. As outlined in **Fig. 1**, the workflow begins with the input query structure – typically the target – and one or multiple subject protein structures in PDB format, all of which can be computational models when an experimental structure is not available.

**Figure 1.**
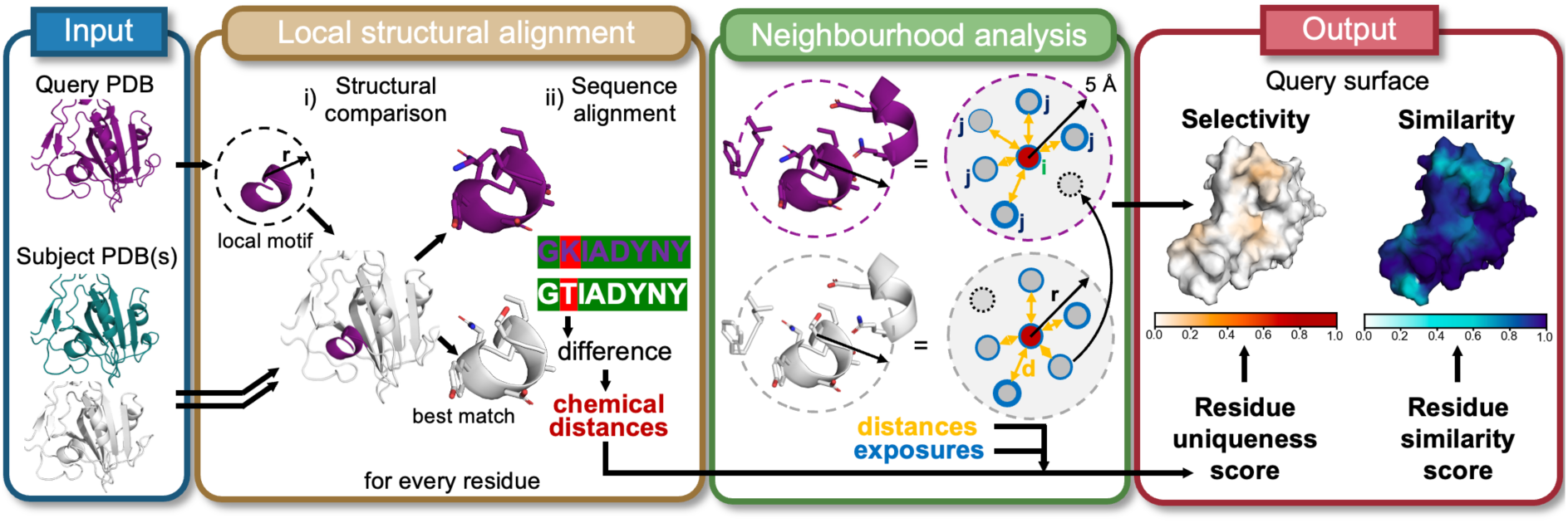
Schematic overview of the SurfDiff pipeline. The workflow begins with the input of query and any number of subject protein structures in PDB format. A local structural alignment module identifies similar motifs based on backbone atom geometry and extract the corresponding sequences to reveal residue-level differences. A subsequent neighbourhood analysis module evaluates the local environment of each residue by computing interatomic distances and relative solvent accessibility (exposure). Residues which are only present in the subject (dotted circle) are added to the query neighbourhood. The final output maps residue-level uniqueness and similarity scores onto the query protein surface, highlighting regions of structural selectivity and conservation.

For each residue in the query protein, the local structural alignment module extracts a structural motif, comprising a central residue with its surrounding residues and uses the MASTER program^32,33^ to search for structurally similar motifs in each subject structure based on backbone atom coordinates. The amino acid sequences of the structurally matching motifs are then retrieved and compared, enabling the identification of identical and different residues, which are characterised in terms of their physicochemical distance (see Methods).

Subsequently, the neighbourhood analysis module defines the local physicochemical environment of each residue by considering all residues within a 5 Å radius. For each central residue, pairwise distances to neighbouring residues within the local surface patch are computed as the minimum atom-to-atom distance. In addition, the relative solvent-accessible surface area (rSASA, see Methods) is calculated to estimate the degree of residue exposure. This analysis is performed across all surface residues. To ensure consistent comparison, residue neighbourhoods are aligned between query and subject proteins. In cases where a residue in the subject has neighbouring residues within 5 Å that are absent in the corresponding query neighbourhood (for example, due to structural divergence or residue substitutions), these neighbours are added to the query neighbourhood with maximal physicochemical distance assigned (see Methods). This supplementation step ensures that all residues present in the subject environment are represented, providing a more complete and sensitive capture of local structural variation.

The final output comprises two scores per surface residue (rSASA > 0.05) mapped onto the query protein surface. Residue uniqueness scores quantify local differences between query and subject structures and serve as indicators of specificity, in the sense that molecules binding to a unique region of the query protein are not expected to bind to any of the subject proteins. Residue similarity scores, by contrast, highlight conserved surface features, aiding in the identification of functional motifs or shared epitopes, so that molecules binding to these regions of the query protein can also be expected to bind all subject proteins. These scores integrate physicochemical, spatial, and exposure-based metrics within each residue’s local context (see Methods).

In simple pairwise comparisons (one query vs one subject), uniqueness and similarity scores are strictly anti-correlated. However, this relationship weakens in one-to-many comparisons, where, for instance, a query motif that resembles one in a subject protein but not in others may score low for both uniqueness and similarity. To enable interpretation at the epitope or domain level, residue-level scores can be aggregated across a defined surface patch, yielding a surface selectivity score (from residue uniqueness) or surface similarity score (from residue similarity). By further integrating these two measures, the surface discriminability score provides a quantitative framework for rational epitope selection, supporting binder design that balances immunogenicity potential (e.g., for antibody or nanobody discovered via immunisation), cross-species conservation, and avoidance of off-target binding. Altogether, SurfDiff offers a sensitive, structure- and sequence-informed scoring method for identifying features relevant to binding specificity, cross-reactivity, and potential immunogenicity, which we demonstrate in the following sections using diverse case studies.

### Surface selectivity scores reflect functional antibody binding differences across SARS-CoV-2 variants

To evaluate the expediency of SurfDiff, we first applied the pipeline to the receptor-binding domain (RBD) of the SARS-CoV-2 spike protein across multiple viral variants. Using the wild-type (WT) Wuhan strain as the query, we calculated residue uniqueness scores through pairwise comparisons with other variants and visualised them on the RBD surface (**Fig. 2A**). For each considered variant, surface selectivity scores were computed over the epitope of the COVOX-40 antibody, originally developed against the WT strain^34^. The resulting scores distinguished between binding and non-binding variants, with the Omicron variant exhibiting the highest surface selectivity score, consistent with pronounced epitope divergence (**Fig. 2B**).

**Figure 2.**
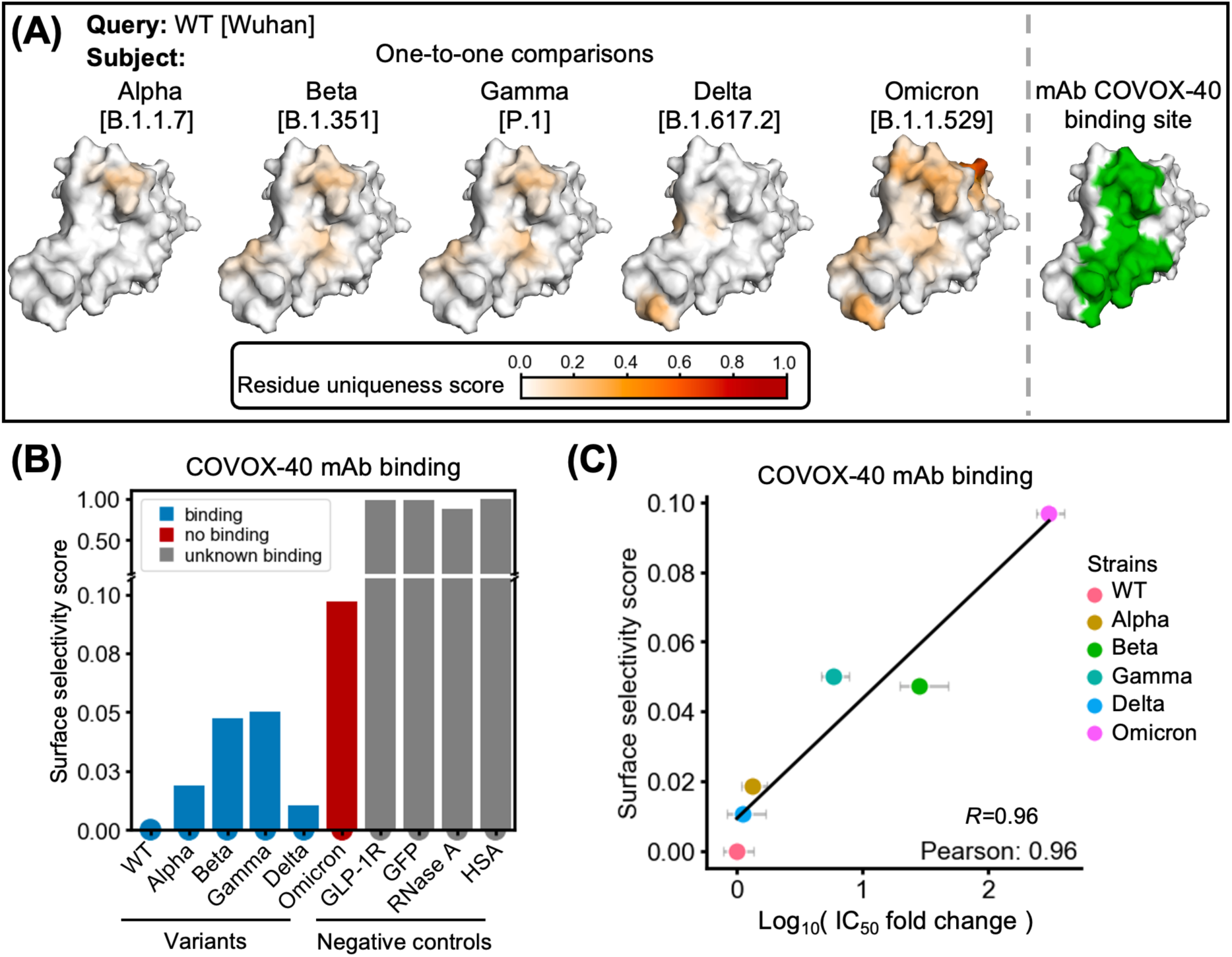
Surface selectivity scores reflect antibody binding differences across SARS-CoV-2 variants. **(A)** Residue uniqueness scores mapped onto the surface of the wild-type (Wuhan) SARS-CoV-2 spike receptor-binding domain. Highly divergent residues (dark red) indicate regions of high structural or sequence variation. The COVOX-40 antibody epitope is outlined in green (PDB: 7nd3). **(B)** Surface selectivity scores of the COVOX-40 epitope computed for multiple SARS-CoV-2 variants (x-axis) using the WT RBD as the query. Structurally unrelated proteins – glucagon-like peptide receptor 1 (GLP-1R; a representative GPCR), green fluorescent protein (GFP; a β-sheet-rich protein), ribonuclease A (RNase A; an α/β-fold protein), and human serum albumin (HSA; a predominantly α-helical extracellular protein) – were included as negative controls. **(C)** Correlation between these surface selectivity scores and experimentally determined inhibitory potency differences, expressed as fold changes in IC₅₀ (that is log₁₀[IC₅₀_variant / IC₅₀_WT]).

To determine whether these scores reflect functionally meaningful differences, we compared them with experimentally determined IC₅₀ values across viral variants^35–39^, expressed as log₁₀-ratios relative to the IC_50_ towards the WT strain (**Fig. 2C**). These transformed values quantify fold differences in inhibitory potency and serve as a proxy for target selectivity – that is, the extent to which a binder discriminates between structurally related targets – rather than absolute binding strength. Notably, SurfDiff-derived surface selectivity scores showed a strong correlation with these potency differences (Pearson *R* = 0.96), despite being computed without any information about the binding molecule itself (including its structure, binding affinity, or chemical properties). Instead, the scores are derived solely from comparative analysis of the binding-region target surfaces, highlighting SurfDiff’s ability to detect intrinsic structural features that underlie differential binding behaviour, independent of any prior knowledge about the interacting molecule.

We further extended the analysis to eight additional monoclonal antibodies with available inhibitory data and epitope information^35–39^. Across all eight antibodies, surface selectivity scores consistently correlated with observed potency shifts (median *R* = 0.88; **Fig. S1**). Finally, we note that equilibrium dissociation constants (K_D_), which directly measure binding affinity, can be employed instead of IC₅₀ in analogous comparisons resulting in very similar correlations (**Fig. S2**). We prioritised IC₅₀ because it is a more functionally relevant readout, and because experimental K_D_ values were not available for as many viral variants.

### Generalisation across diverse binder types

Encouraged by the results on antibody-RBD complexes, we applied the method to a representative set of small molecules, peptides, and antibodies targeting a range of unrelated proteins to evaluate the applicability of SurfDiff across distinct molecular recognition contexts. Each binder had one known co-crystal structure and experimentally determined binding affinities to a range of targets (expressed as K_i_ or K_D_ values)^40^. In each case, surface selectivity scores were computed across a panel of related proteins or protein variants (the targets) and compared to log10 fold-changes in affinity from experiments.

SurfDiff scores calculated solely from the protein targets showed strong Pearson correlations with affinity-derived selectivity values across all binder and target classes, indicating that the method generalises effectively across different classes of molecular binders and target proteins. For small-molecule ligands – including endogenous serotonin, 5-carboxamidotryptamine, and the antiparkinson 5-HT2A agonist lisuride – strong correlations were observed across serotonin receptor subtypes, with lisuride additionally showing consistent selectivity patterns across distinct GPCR families (**Fig. 3A-C**). Peptide ligands such as vasopressin, oxytocin, and somatostatin-14 also demonstrated robust correlations between surface selectivity scores and experimentally determined affinity differences within their respective receptor families (**Fig. 3D-F**). Similarly, therapeutic antibodies – including 4E11 (targeting dengue virus envelope protein DIII)^41^, N6 (targeting HIV-1 gp120 protein)^42^, and fresolimumab (targeting TGF-β family cytokines)^43^ – exhibited the expected trend between SurfDiff scores and measured selectivity differences across antigen variants and isoforms, albeit only 3 or 4 experimentally determined data points were available for these antibodies (**Fig. 3G-I**).

**Figure 3.**
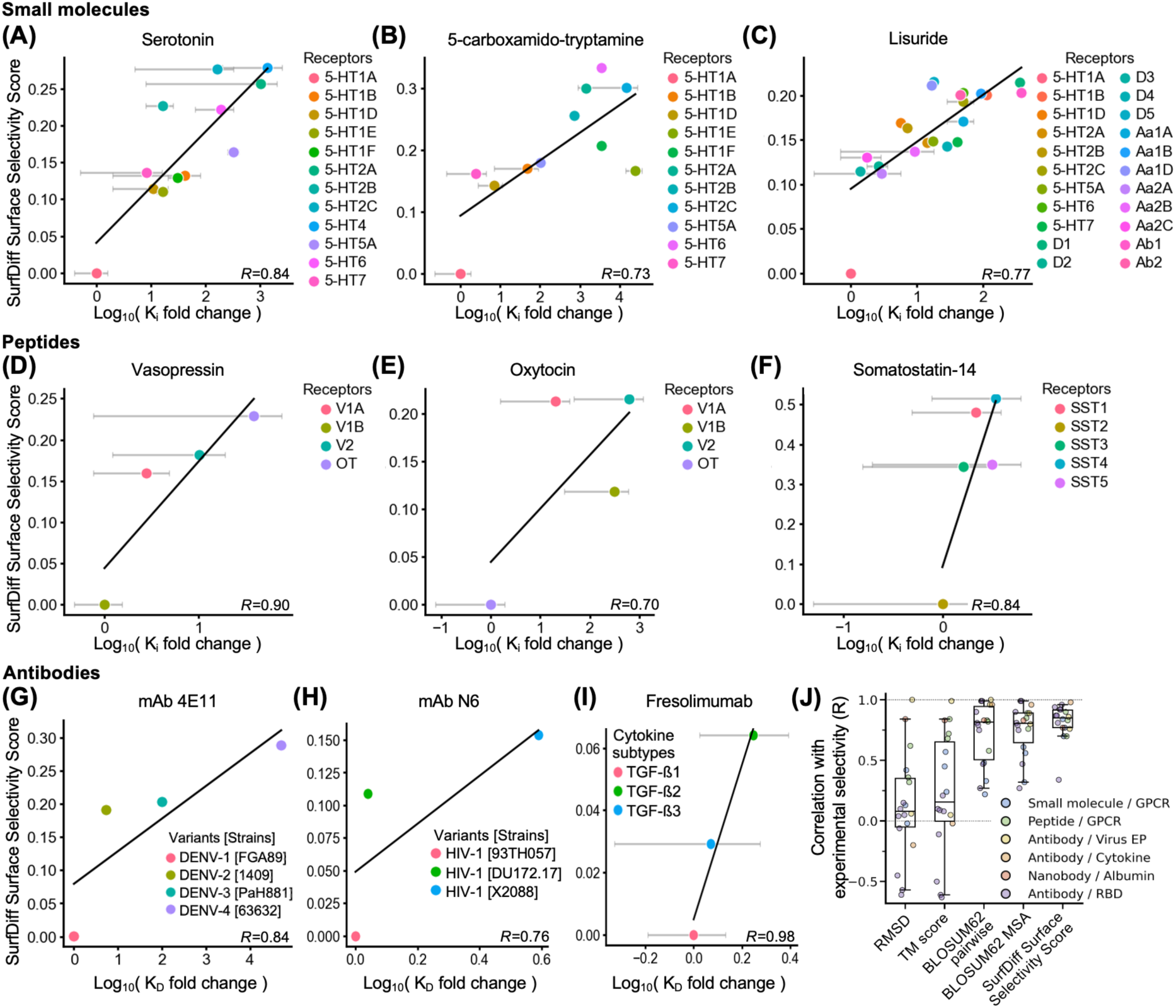
Generalisation of surface selectivity scores across diverse binder types. (A-C) Correlation between SurfDiff surface selectivity scores and experimentally determined binding selectivity (expressed as Log_10_ fold changes in K_i_ or K_D_ values) for small-molecule ligands: **(A)** Serotonin binding across serotonin GPCR subtypes (legend); **(B)** 5-carboxamido-tryptamine binding across serotonin GPCR subtypes; **(C)** Lisuride binding across serotonin, dopamine, and adrenergic GPCR subtypes. **(D–F)** Peptide–receptor interactions: Vasopressin **(D)** and Oxytocin **(E)** binding across vasopressin and oxytocin receptor subtypes; **(F)** Somatostatin-14 binding across somatostatin receptor subtypes. **(G–I)** Antibody-antigen systems: **(G)** Monoclonal antibody 4E11 binding to envelope protein DIII from multiple dengue virus strains; **(H)** Broadly neutralising antibody N6 binding to HIV-1 envelope gp120 protein; **(I)** Therapeutic antibody Fresolimumab binding to members of the TGF-β cytokine family. In all panels, the *y*-axis represents SurfDiff surface selectivity scores, while the *x*-axis shows Log₁₀-transformed fold differences in K_i_ or K_D_ relative to the query target. Unless otherwise specified, K_i_ values were obtained from the IUPHAR Guide to Pharmacology database. Where the original studies reported ranges instead of mean and standard error, the mean values between reported minima and maxima were used for correlation analysis (as data point and error bar ranges, respectively). Pearson correlation coefficients (*R*) are indicated for each plot. **(J)** Extended analysis across all binder-target-variants systems considered, reported as boxplot of Pearson correlation coefficients between experimentally determined selectivity (i.e., inhibitory potency or binding fold-changes), and various metrics calculated from epitope residues (x-axis). For metrics for which an anticorrelation was expected (TM score and BLOSUM62-based metrics), the sign of the Pearson’s coefficients was changed to allow for direct visual comparison of correlation strengths across metrics. Boxes represent the first and third quartiles of the distribution, whiskers represent the 1.5 interquartile range, the median is the horizontal bar. A Friedman’s test yielded P-value = 3×10^-5^, N = 18.

For comparison, we conducted analogous analyses on all classes of binders and targets by employing, in place of the SurfDiff surface selectivity score, commonly used metrics calculated from the epitope residues. Specifically, we tested the epitope backbone root mean square deviation (RMSD), its structural TM-score, and BLOSUM62 substitution scores calculated from differences in epitope residues, both by doing individual pairwise alignments between the WT and each variant sequence under scrutiny, or by doing a single multiple-sequence alignment (MSA) encompassing WT and all variants (see Methods). We find that SurfDiff outperforms all these established metrics, by yielding consistently stronger correlations with experimentally observed selectivity (**Fig. 3J**).

Collectively, these results demonstrate that SurfDiff effectively captures surface-driven selectivity determinants across a wide range of binder classes and target families, independent of the nature of the binding molecule.

### Characterising Subtype-Selective Binding Sites in Structurally Conserved GPCRs Using Pairwise Surface Comparison

To further illustrate the versatility and practical utility of SurfDiff, we applied it to six representative case studies spanning diverse biomedical research areas. In each instance, similarity and difference analyses were directly compared to experimental binding data from characterised binders, thus validating the calculated selectivity profiles.

Subtype-selective targeting remains a significant challenge in drug development for closely related receptor or pathogenic-protein families, where high structural conservation often masks functionally important differences^26^. To demonstrate how SurfDiff can help resolve such subtleties, we analysed angiotensin II receptor subtypes type 1 (ATR1) and type 2 (ATR2) – two closely related GPCRs activated by the same endogenous ligand, angiotensin II (AngII), but eliciting opposing physiological effects relevant in blood pressure regulation^44,45^. Despite their high structural similarity, ATR1 and ATR2 represent distinct therapeutic targets requiring selective modulation.

We applied SurfDiff to identify subtype-specific surface divergence between ATR1 and ATR2, focusing on two key regions: the shared AngII binding site and the ATR2-specific epitope recognised by the Fab 4A03^46^. To benchmark SurfDiff’s sensitivity, we also compared ATR2 with glucagon-like peptide receptor 1 (GLP-1R), a peptide-binding GPCR of similar topology but unrelated function. **Fig. 4A** presents structural models of ATR1, ATR2, and GLP-1R, with mapped residue uniqueness scores highlighting local surface divergences.

**Figure 4.**
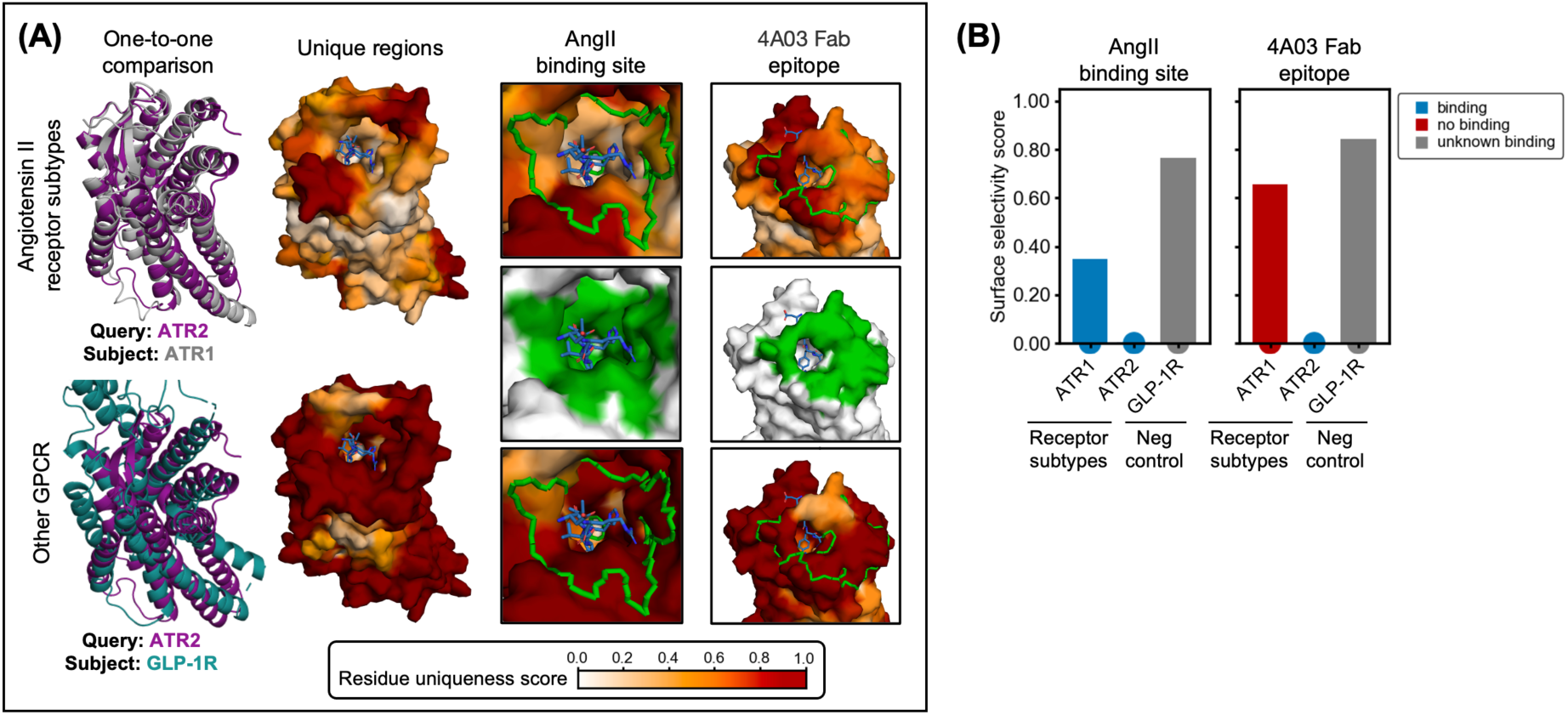
Surface selectivity analysis of angiotensin II receptor subtypes ATR1 and ATR2. **(A)** Structures of ATR1 (PDB: 6os2), ATR2 (PDB: 6jod), and glucagon-like peptide receptor 1 (GLP-1R) (PDB: 6×18). Residue uniqueness scores are mapped onto the surface of ATR2. The angiotensin II (AngII) binding site and the epitope of the ATR2-specific Fab 4A03 are highlighted in green. ATR2 is compared with GLP-1R serving as a structurally related negative control. **(B)** Bar plot of surface selectivity scores for the AngII binding site and 4A03 Fab epitope.

Consistent with the conserved binding affinity of AngII to both ATR1 and ATR2 (K_i_ 0.2 nM and 0.3 nM, respectively^47^), the ligand-binding site displays comparably low surface selectivity scores (**Fig. 4B**), reflecting reasonable conservation. By contrast, the Fab 4A03 epitope exhibits elevated surface selectivity scores, in line with its reported ATR2-specific binding^46^. Importantly, comparison of ATR2 with the homologous GLP-1R receptor reveals extensive surface divergence across both sites, underscoring SurfDiff’s ability to discriminate among topologically similar peptide-binding GPCRs.

These results, together with those in **Fig. 3**, demonstrate that SurfDiff can sensitively resolve subtype-selective surface features within GPCR families, distinguishing conserved and divergent patches, an information that is crucial for guiding the design of selective therapeutic binders.

### Mapping Conserved and Divergent Epitopes Across Isoforms and Viral Variants via Multi-Target Surface Profiling

SurfDiff facilitates systematic comparison of surface features across multiple related protein targets, offering insights into both conserved and subtype-specific regions critical for the rational design of selective binders and useful to inform immunogen engineering. Extending beyond pairwise comparisons, this one-to-many approach reveals patterns of local surface similarity and divergence across entire protein families or homologous variants.

We applied this methodology to two biologically and therapeutically relevant examples: the transforming growth factor-beta (TGF-β) cytokine isoforms^48^ and the HIV envelope protein gp120 from diverse viral variants^7^. Both cases demonstrate SurfDiff’s utility in pinpointing conserved epitopes suitable for pan-subtype targeting, alongside unique surface patches that may serve as foundations for subtype-specific binder development, for example for diagnostic purposes.

For the TGF-β family, we compared surface features among the three closely related isoforms (TGF-β1, TGF-β2, TGF-β3) using a one-to-two comparison scheme, whereby each isoform was evaluated against the surfaces of the other two (**Fig. 5A**). This analysis highlighted broadly conserved regions as well as isoform-specific surface patches. The epitope of the pan-subtype antibody fresolimumab, which binds all three isoforms with high affinity (1-3 nM across all isoforms)^43^, was examined in detail. Consistent with its broad binding profile, high similarity (**Fig. 5A**) and low surface selectivity scores were observed at the fresolimumab epitope (**Fig. 5B**), reflecting strong local conservation. In contrast, outside of this epitope, elevated residue uniqueness scores reveal isoform-specific patches that may be exploited for selective binding (**Fig. 5A**).

**Figure 5:**
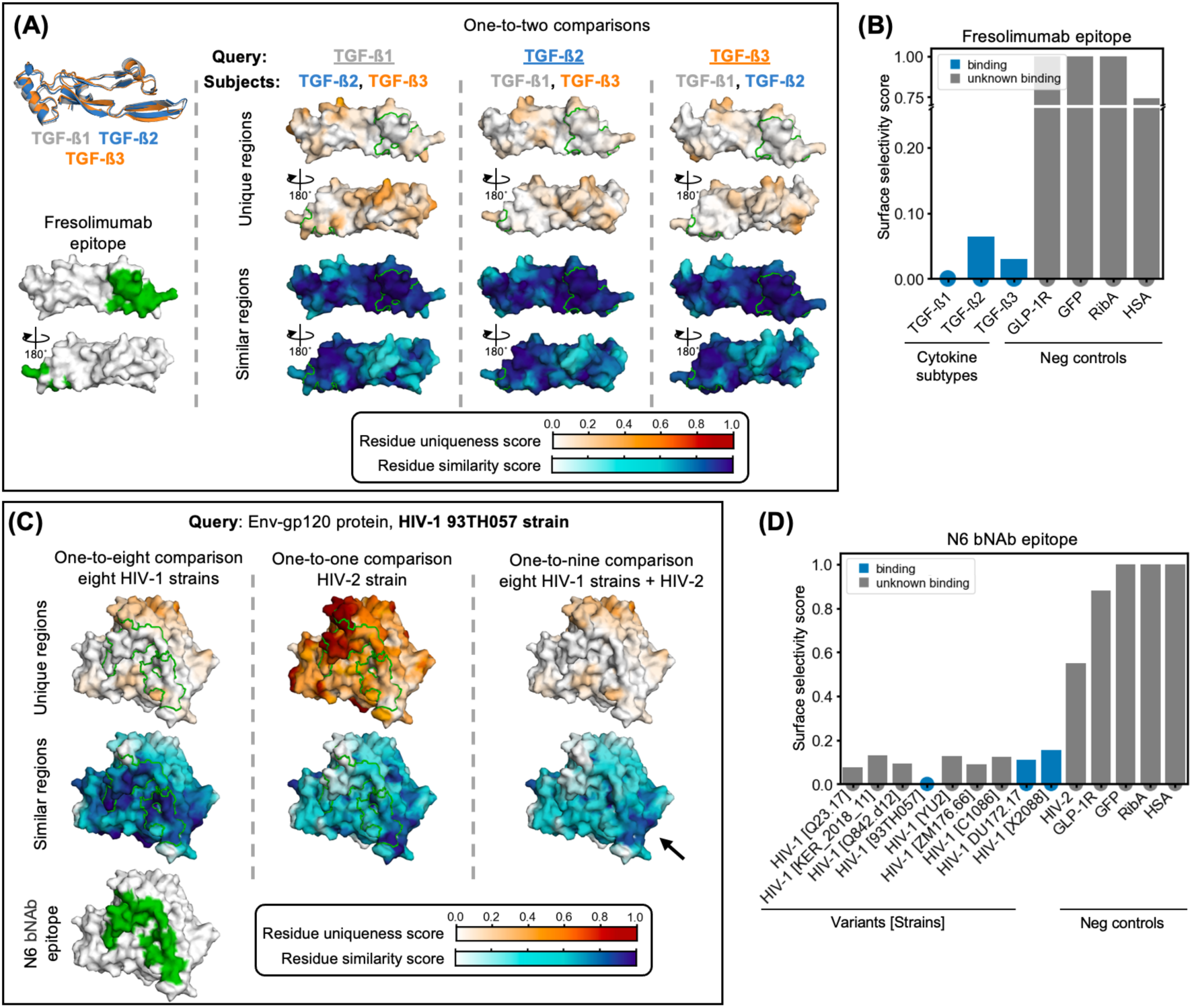
Mapping conserved and subtype-specific surface features across related proteins using SurfDiff. **(A)** Residue similarity and uniqueness scores mapped onto surfaces of the three human TGF-β isoforms (TGF-β1, TGF-β2, TGF-β3; PDB: 4kv5, 4kxz, 3eo1). Each isoform was compared to the other two in a one-to-two fashion. Regions of local surface divergence (high uniqueness) and conservation (high similarity) are highlighted. The epitope of the pan-subtype antibody fresolimumab is bordered in green. **(B)** Bar plot of the surface selectivity scores for the fresolimumab epitope. **(C)** Residue similarity and uniqueness scores mapped onto the surface of the HIV-1 gp120 reference variant (strain: 93TH057, PDB: 5te6), using three comparison sets: (i) against eight other HIV-1 variants, (ii) against one HIV-2 variant, and (iii) against all nine variants combined (HIV-1 + HIV-2). The broadly neutralizing antibody (bNAb) N6 epitope is bordered in green. The combined comparison highlights a small number of surface patches with moderate to high similarity across all variants, representing potential targets for pan-HIV-1/2 neutralization strategies (black arrow). **(D)** Bar plot of the N6 epitope surface selectivity scores from 1:1 comparisons (x-axis). This score is low in intra-HIV-1 comparisons reflecting broad conservation and high against HIV-2, indicating limited or no cross-reactivity.

In the HIV example, we focused on the gp120 envelope protein, a key viral target whose extensive variability challenges broad neutralisation^7^. Using a well-characterised HIV-1 reference variant, we conducted three complementary comparisons: (i) against eight other HIV-1 variants to assess intra-variant conservation; (ii) against a single HIV-2 variant to evaluate inter-variant divergence; and (iii) against all nine variants combined to integrate both levels of variability. In particular, we focussed on the epitope of the broadly neutralising antibody (bNAb) N6, which neutralises HIV-1 variants, but was not reported to bind to HIV-2^42^.

As anticipated, N6’s epitope corresponded to a highly conserved surface patch among HIV-1 variants, indicated by high surface similarity scores (**Fig. 5C**) and correspondingly low surface selectivity scores (**Fig. 5D**). Conversely, comparison with HIV-2 revealed marked divergence at this site, consistent with known dissimilarity between HIV-1 and HIV-2^49^. The combined HIV-1 versus HIV-1 + HIV-2 analysis revealed a layered pattern: residue uniqueness scores predominantly reflected intra-HIV-1 variability, while residue similarity scores identified a limited set of surface patches with moderate to high similarity across all variants (**Fig. 5C**), highlighting candidate regions for the potential development of truly pan-HIV-1/2 antibodies (black arrow in **Fig. 5C**). We note, however, that other aspects that are important for binding, such as glycosylation patterns, are not considered in this analysis, which is simply meant as an example application.

Together, these examples illustrate how SurfDiff’s residue similarity and uniqueness scores offer complementary perspectives: residue similarity identifies broadly conserved regions suitable for pan-subtype binder design, whereas residue uniqueness highlights divergent sites ideal for subtype-specific targeting. This dual insight is essential for rational immunotherapeutic design, enabling tailored development of binders with either broad cross-reactivity or high selectivity, depending on the desired biological and clinical objectives.

### Cross-species epitope prioritisation using the surface discriminability score

Rational epitope selection underpins the design of binding therapeutic candidates across modalities, including antibodies, nanobodies, engineered proteins and peptides. It must balance discovery constraints (e.g., immunogenicity in host species when immunisation is used), the species cross-reactivity required for preclinical pharmacology and toxicology, avoidance of off-target interactions, and, where appropriate, broad neutralisation or pan-haplotype binding. To address these competing requirements in a binder-agnostic manner, we introduce the surface discriminability score, a residue-level metric that integrates residue uniqueness and similarity across prespecified target and non-target sets, and quantifies the trade-off between breadth across desired targets and exclusion of undesired off-targets. We applied this framework to systems with well-characterised experimental datasets: serum albumin **Fig. 6A-C**) and the immune checkpoint receptor programmed cell death protein 1 (PD-1) (**Fig. 6D-F**).

**Figure 6:**
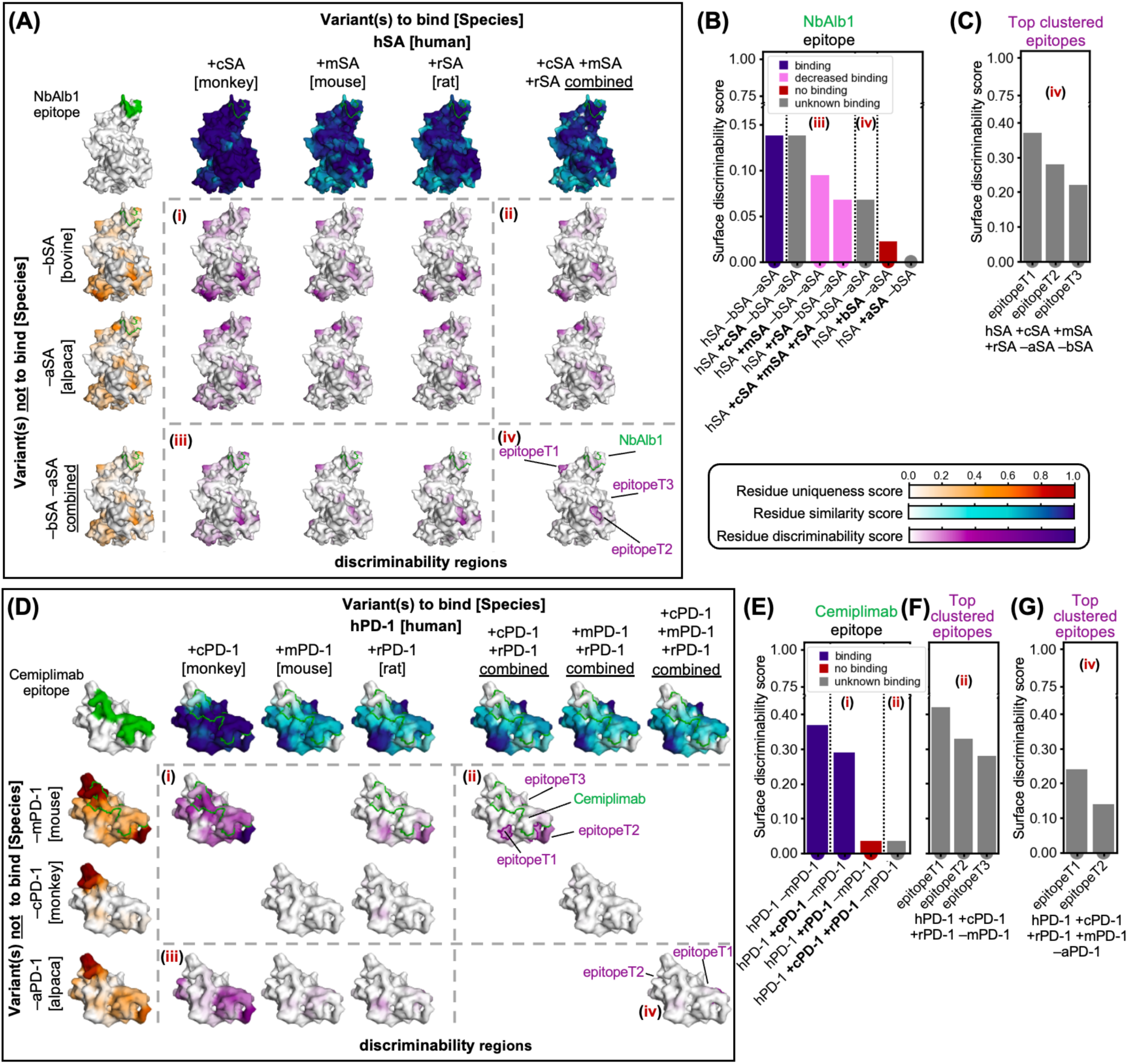
Cross-species epitope prioritisation with the discriminability score. (**A**) The header row shows residue similarity scores mapped onto the surfaces of human serum albumin (hSA), used as query, compared to orthologues from cynomolgus monkey (cSA), mouse (mSA), rat (rSA) used as subjects (i.e., variants to bind), while the header column shows hSA residue uniqueness scores compared to alpaca (aSA) and bovine serum albumin (bSA) used as subjects (i.e., variants not to bind). The epitope of the camelid-derived nanobody NbAlb1 is highlighted in green. The matrix (within dashed lines) shows discriminability scores mapped on hSA for all possible comparisons, in block (i) there are simpler surface comparisons (e.g., bind hSA +cSA and not –bSA), while (ii), (iii), and (iv) have more complex comparisons (for example iv corresponds to hSA +cSA +mSA +rSA –bSA –aSA). The top 3 clustered epitopes across all species are labelled on (iv), and highlight regions with high surface discriminability scores, that is conserved across variants to bind and distant to variants not to bind. (**B**) Bar plot of the surface discriminability scores at the NbAlb1 epitope, all calculated by requiring not to bind aSA and bSA. The score is high for hSA, low for bSA, and intermediate for mSA and rSA, consistent with experimentally observed binding patterns (legend). (**C**) SurfDiff-predicted top 3 clustered epitopes on hSA (iv) with elevated surface discriminability scores across hSA, cSA, mSA, and rSA, providing candidate sites for broad yet selective albumin recognition while avoiding bSA and aSA. (**D**) Like **A**, but for human PD-1 (hPD-1) and orthologues from cynomolgus monkey (cPD-1), mouse (mPD-1), and rat (rPD-1), with alpaca PD-1 (aPD-1) included to assess suitability for immunisation to raise nanobodies, and mouse and monkey considered both as targets for species cross-reactivity and as a potential immunisation hosts (not to bind). (**E**) Surface discriminability scores at the Cemiplimab epitope: high for hPD-1 –mPD-1, (i) high for hPD-1 +cPD-1 –mPD-1, low for hPD-1+rPD-1 –mPD-1, and low for hPD-1 –mPD-1 reflecting known binding patterns (legend); (**F**) additional SurfDiff-predicted top 3 epitope clusters with elevated scores, including regions supporting rPD-1 binding while maintaining cPD-1 binding. (**G**) Top cross-species epitope clusters (iv) across hPD-1, cPD-1, mPD-1, and rPD-1, highlighting candidate sites for generating broadly cross-reactive nanobodies, while confirming sufficient divergence from aPD-1 for immunisation.

Albumin binding is a key determinant of pharmacokinetics and can markedly extend half-life, especially for smaller proteins^50^. For example, anti-human serum albumin (hSA) nanobodies are often fused to bioactive targeting nanobodies to increase exposure^51^. Therefore, therapeutic candidates must ideally recognise serum albumin (SA) of common preclinical species, such as cynomolgus monkeys (cSA), mouse (mSA), and rat (rSA), while avoiding bovine SA (bSA), which is widely used as a blocking reagent in *in vitro* and *ex vivo* assays. Moreover, the epitopes of nanobodies discovered by immunisation would typically retain sufficient divergence from camelid SA (e.g., alpaca; aSA) to elicit a response (**Fig. 6A**). The camelid-derived nanobody NbAlb1 was selected, following immunisation, via laboratory screening against multiple SA variants and hence exemplifies this balance: it binds hSA but not bSA and shows reduced affinity for mSA and rSA^52–54^, a pattern mirrored by the discriminability scores for its epitope (**Fig. 6B**). In addition to NbAlb1, **Fig. S3** reports equivalent analyses for six further hSA binders, including nanobody Nb.b201 (discovered from a synthetic library via yeast display)^55^, Fab CA645^56^, shark VNAR huE06v1.1^57^, an affibody^58^, and the human neonatal Fc receptor (FcRn)^59^. Beyond these already characterised epitopes, SurfDiff identified epitope clusters with even higher discriminability across hSA, cSA, mSA and rSA while remaining distinct from bSA and aSA (**Fig. 6A(iv), Fig. 6C, Tab. S1**), pointing to promising sites for broad yet selective albumin recognition.

For human PD-1 (hPD-1), preclinical studies similarly require antibodies that engage hPD-1 together with one or more orthologues (cPD-1, mPD-1, rPD-1). Cemiplimab, an FDA-approved antibody^60^ generated by immunising transgenic mice with hPD-1^61^, binds hPD-1 and cPD-1 but not rodent orthologues. Although these mice are genetically humanised (yielding human antibody sequences), immune tolerance can limit responses to self-antigens such as mPD-1. This pattern is reflected in discriminability scores for the Cemiplimab epitope, which are high for hPD-1 and cPD-1 but low for mPD-1 and rPD-1 (**Fig. 6E**). Beyond this known epitope, SurfDiff identified additional epitope clusters with similarly high scores, including regions with potential to support rPD-1 binding (**Fig. 6F, Tab. S2**). Comparative analysis of human and alpaca PD-1 indicated sufficient divergence for effective immunisation to discover anti-hPD-1 nanobodies, and SurfDiff highlighted clusters with elevated scores across hPD-1, cPD-1, mPD-1 and rPD-1, offering candidate epitopes for broad-spectrum nanobody generation (**Fig. 6D(iv), Fig. 6G, Tab. S3**). An additional example involving human transferrin receptor 1 (TFR-1), a common target for shuttling biologics across the blood–brain barrier^62^, is presented in **Supplementary Fig. S4**.

Taken together, these analyses illustrate that SurfDiff discriminability provides a binder-agnostic, modality-independent criterion for cross-species epitope prioritisation, compatible with discovery by immunisation, *in vitro* selection, or computational design.

## Discussion

Conventional drug or affinity-reagent discovery pipelines typically yield binders with little or no control over where they bind. Functional or competition assays, when used for screening, can enrich for a region of interest, but they seldom define the binding footprint at single-residue resolution. Experimental epitope mapping for discovered hits can be as or more labour-intensive than direct functional characterisation of hits across all desired targets and undesired off-targets. However, this landscape is rapidly changing. Accurate *in silico* epitope mapping from structural modelling, together with de novo binder design aimed at predetermined epitopes of interest, are increasingly making epitope choice a prospective design variable rather than a retrospective outcome.

In this context, this study introduces SurfDiff, a structure- and sequence-informed framework for residue-level comparison of protein surfaces that integrates local structural alignment with physicochemically and spatially informed neighbourhood analysis. Designed to capture subtle surface variations that govern binding specificity, cross-reactivity, and immunogenic potential, SurfDiff provides a generalisable approach applicable across diverse protein families and interaction types, ranging from viral antigens and serum albumin to cytokine isoforms and GPCR subtypes.

Similarly, recent progress in structural vaccinology – accurate complex modelling^11,24,25^, escape/antigenic-mapping data^63,64^, and increasingly reliable structural design of immunogens^65–67^ – are making epitope-focused strategies increasingly feasible. What remains largely unaddressed is a prospective, quantitative way of choosing which B-cell epitopes to prioritise across variant lineages while defocusing variable or host-mimicking surfaces. On finding targetable B-cell epitopes on protein surfaces, SurfDiff comparisons can rank conserved epitopes across variant sets, help identify state-specific sites (e.g., prefusion over postfusion), and pinpoint where immunofocusing interventions (e.g., glycan masking, scaffolded or nanoparticle display, mosaic designs, etc.) are likely to be most effective. However, it is not likely useful to model T-cell epitopes (which involve non-surface peptides), germinal-centre dynamics, or glycan heterogeneity.

Central to SurfDiff are two complementary residue-level metrics: residue uniqueness, which highlights divergent surface patches ideal for selective targeting, and residue similarity, which identifies conserved regions enabling cross-reactive binding. Then, the surface discriminability score combines both measures to prioritise epitopes that balance conservation across desired targets with divergence from off-target proteins or undesired species orthologues. Across the hSA and PD-1 case studies it recapitulated known cross-species binding patterns and highlighted additional epitope clusters with favourable profiles. Taken together, SurfDiff scores offer an interpretable, binder-agnostic, quantitative measure for rational epitope prioritisation, thus helping focus discovery or design efforts on epitopes most likely to deliver the desired selectivity profile, while reducing reliance on extensive, unfocused library screenings.

SurfDiff accommodates both one-to-one and one-to-many comparisons. Pairwise analyses, such as ATR1 versus ATR2 (**Fig. 4**), reveal subtle but functionally relevant surface differences critical for subtype-specific targeting while minimising off-target effects. Broader one-to-many comparisons, for example across TGF-β isoforms or HIV gp120 variants (**Fig. 5**), systematically uncover conserved and divergent regions consistent with known patterns of cross-reactivity and strain specificity. This flexibility enables applications ranging from immunogen design to epitope prioritisation in drug discovery.

Validation across diverse systems, including viral antigens, cytokine isoforms, GPCR subtypes, and serum albumin, demonstrated strong concordance with experimental binding data for multiple binder types, including small molecules, peptides, antibodies, and nanobodies. SurfDiff scores correlated closely with relative differences in IC₅₀, Kᴅ, and Kᵢ measurements, confirming that comparative surface features alone capture determinants of relative binding preferences. It is important to note, however, that SurfDiff is not a general predictor of binding affinity, since binding strength depends on the joint contribution of both epitope and paratope (i.e., target and binder), and the method incorporates no information on the binding partner. It focuses on surface divergence and conservation, providing actionable insights for epitope prioritisation, design of selective or cross-reactive binders, mapping conserved regions, and assessing cross-species immunogenic potential. The correlations observed consistently with experimental data underscore the practical value of SurfDiff for these tasks.

Benchmarking against standard approaches to compare epitopes, such as backbone RMSD, TM-score, and BLOSUM62 similarity confirmed that SurfDiff captures functionally relevant surface differences with much higher sensitivity, by integrating physicochemical, spatial, and exposure information. Moreover, tests with alternative PDB entries showed robustness to input model variation (**Tab. S5**), an encouraging feature given the growing use of predicted structures such as AlphaFold models when experimental structures are unavailable.

SurfDiff operates on local surface geometry and chemistry and is applicable to experimental structures and predicted models, without requiring high global structural similarity. It can also be applied to distantly related – or even unrelated – proteins (as in our negative-control comparisons), where scores remain interpretable, although systematic benchmarking is limited by the lack of ground-truth data on cross-reactivity in this regime. Because SurfDiff is inherently comparative, performance depends on the choice of query and subject structures. Inappropriate or low-quality structures or models, or including single conformations of proteins that populate multiple ones, will affect accuracy. We expect that the reliability of SurfDiff will degrade when applied to sites that are highly dynamic (e.g., intrinsically disordered regions or flexible loops). For linear, flexible epitopes, multiple-sequence alignments or motif-conservation analyses may suffice and indeed even outperform SurfDiff when dynamic is very high. For conformational, state-dependent epitopes, analysing ensembles spanning functional states (open/closed, active/inactive, etc.) can help identify state-specific features. Future extensions will include explicit treatment of glycosylation and other post-translational modifications, membrane context and condition-specific accessibility, thus further refining the analysis of complex biomolecular systems.

In summary, SurfDiff provides a versatile and interpretable framework for single-residue-resolution protein surface comparison, detecting both conserved and divergent motifs across related and unrelated proteins. By integrating physicochemical, spatial, and solvent-exposure information, it supports applications such as epitope mapping, immunogen design, and rational development of selective or cross-reactive binders. The surface discriminability score further enhances this approach by enabling quantitative epitope prioritisation that balances potential immunogenicity, preclinical relevance, and avoidance of off-target interactions, providing a predictive and translationally relevant platform for therapeutic and diagnostic applications.

## Materials and Methods

### Overview of the Pipeline

The SurfDiff pipeline is designed to compare local patches of protein surfaces at the residue level, integrating structural, sequence, and physicochemical information to identify conserved or divergent epitopes/binding sites. In brief, the workflow consists of the following steps: (i) definition of local structural motifs for each residue; (ii) exhaustive structural alignment of these motifs across query and subject proteins; (iii) sequence comparison and physicochemical characterisation of matched motifs; (iv) definition of local neighbourhoods and calculation of residue-level uniqueness (RUS), similarity (RSS), and, if asked, discriminability scores (RDS); (v) aggregation of residue-level scores across surface patches to quantify regional conservation, uniqueness, or cross-species/variant reactivity and discriminability. The pipeline supports both one-to-one comparisons, in which a single query is compared to a single subject, and one-to-many comparisons, where a single query is compared to multiple subjects to estimate conserved or divergent features.

### Structural Data Acquisition and Preparation

Protein structures used as input for the SurfDiff pipeline are provided in PDB format. These can be either experimentally determined structures or high-confidence computational models. For this study, structures were either obtained from the Protein Data Bank^68^ or modelled from their Uniprot sequence entry^69^ using the Alphafold3 webserver (https://alphafoldserver.com)^11^. All used PDBs and modelled structures are provided in **Tab. S5**.

Prior to analysis, all structures were standardised using in-house PDBcleaner software available on our webserver (https://www-cohsoftware.ch.cam.ac.uk/index.php/pdbcleaner). This included selection of relevant protein chains and removal of all non-protein atoms such as water molecules, ions, and ligands. Missing sidechain atoms were modelled.

### One-to-one Comparisons: Definition and Alignment of Local Structural Motifs

For each residue in the query protein, a local structural motif is defined to capture its immediate three-dimensional context. Each motif comprises the central residue and its neighbouring residues, identified based on the proximity of their Cα atoms within a radius of 3 to 7 Å. These residue-centric motifs form the basis for detailed local comparisons between query and subject structures.

Structural alignments are performed using the MASTER algorithm version 1.6^32,33^, which enables exhaustive and efficient comparison of each query motif, including discontinuous tertiary motifs, against all potential substructures in the subject protein. ‘Hit’ motifs are defined as those with backbone RMSD ≤ 2.0 Å, and these are sorted by increasing RMSD values.

To reduce coincidental matches from repetitive or highly regular structural elements, such as α-helices, a discrimination score is introduced. In backbone-only comparisons, structurally similar motifs can yield similarly low RMSD values, particularly in proteins dominated by repetitive elements, potentially aligning more closely to unrelated but geometrically similar regions than to true homologous counterparts. The discrimination score, defined as *(RMSD_best_ – RMSD_second-best_)/motif_lenght*, assesses the reliability of the top-ranking hit (within a given subject) by measuring how much its RMSD exceeds that of the second-best hit, normalized by the motif length. A threshold of ≥ 0.005 Å/residue is used to accept the top hit as a reliable structural correspondence.

If no motif meets these criteria, the residue is temporarily labelled as different. Matching is then repeated using progressively larger motifs by incrementally expanding the radius around the central Cα atom to 4, 5, 6, and 7 Å until the criteria are met (at which point the sequence comparison is carried out). Residues failing to match after the final round are classified as different.

### One-to-one Comparisons: Sequence Comparison and Physicochemical Characterisation

Once a first confident structural match is identified, the amino acid sequence of the aligned motif in the subject protein is extracted and compared to the corresponding query motif. Sequences are aligned solely based on the local structural superimposition. Pairwise comparisons identify identical residues and substitutions. A matched motif is considered sufficiently similar if it contains three or more identical residues. If this criterion is not met, the structural alignment and sequence comparison are repeated by expanding the query motif is incrementally (4 - 7 Å like before). Residues failing to meet the threshold after the final expansion are classified as different.

For each query residue, the sequence of the matched motif that satisfies both structural and sequence similarity thresholds is collected into a positional grid resembling a multiple sequence alignment. Each row corresponds to the short sequence of a matched subject motif, which may be discontinuous, as sequence-distant fragments can occupy the same structural motif in 3D space. The most frequently occurring residue at each position across all matched motifs is selected as the consensus residue, serving as the reference for defining the physicochemical distance to the query residue.

Residues for which no matching subject motif was identified are assigned the maximum score of 1, as these query residues have no corresponding subject residue and are considered maximally different. For all others, residue substitutions are scored using a modified BLOSUM62 substitution matrix^70^ representing physicochemical distance, with diagonal values (self-matches) set to zero to isolate non-identical substitutions. Scores are min–max normalised to range between 0 and 1, reflecting physicochemical differences rather than evolutionary background frequencies (**Tab. S4**).

### Neighbourhood Definition and Local Environment Analysis

The local chemical environment of each residue *i* in the query (𝑄) comprises all neighbouring residues *j* with at least one heavy atom within 5 Å of any atom of *i*. The same procedure is applied to subject (*S*) residues to ensure consistency. Formally, *j* is a neighbour of *i* if:

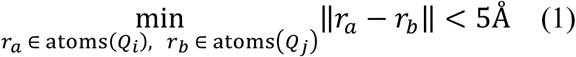

where *r_a_* and *r_b_* are the 3D Cartesian coordinates of atoms *a* in residue *i* and *b* in residue *j*, respectively. These neighbourhoods provide the local surface context and are subsequently used to compute spatial features.

The minimum interatomic distance between each central–neighbour pair (*i*,*j*) is stored as a feature of local geometry:

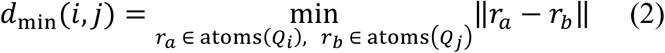

For consistent comparison, each query residue *Q_i_* is associated with its consensus-matched residue *S_i_* in the subject. If a neighbourhood residue *S_j_* is present in the subject but absent in the query, it is added to the query neighbourhood with the maximum physicochemical distance. This ensures that all subject residues contribute to the local patch characterisation.

### Calculation of Residue-Level Uniqueness, Similarity and discriminability Scores in one-to-one comparisons

Residue *i* in the query is assigned two complementary scores describing its local structural and chemical relationship to the corresponding residues *j* in the subject:

The residue uniqueness score of residue *i* (RUS*_i_*) quantifies divergence between query and subject by combining physicochemical dissimilarity (𝑑_*chem*_) with spatial (𝑤_*dist*_, Eq. 4) and solvent exposure (𝑤_*exp*_;<, Eq. 5) weights:

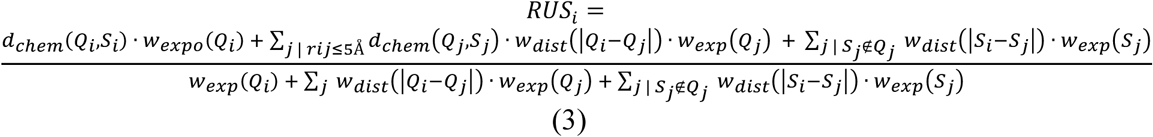

Where *Q* and *S* stand respectively for query and subject residue. For a central-neighbour residue pair Q*_i_*,Q*_j_* (or, analogously, S_i_,S_j_), the minimum interatomic distance is denoted as 𝑥 = |𝑄_i_ – 𝑄_j_| = 𝑑_min_(𝑖, 𝑗). Then, the distance-based weighting function 𝑤*_dist_*:(𝑥) (Eq. 4) assigns full weight to neighbours within 1 Å, decreases linearly to 0 at 5 Å, and excludes any residue beyond 5 Å:

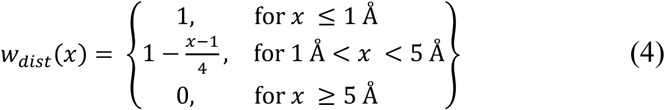

Relative solvent-accessible surface area (rSASA) is computed for each residue by calculating SASA using the Shrake-Rupley algorithm with a 1.4 Å probe radius and dividing it by the SASA of the same amino acid (X) in an extended Gly-X-Gly tripeptide. Then, solvent exposure weights 𝑤*_expo_*(𝑥) (Eq. 5) are calculated using a modified Hill function, with a threshold set at *x* = rSASA_i_ = 0.05, effectively down-weighting buried residues.

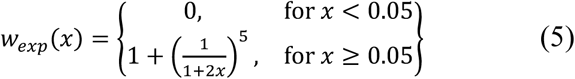

Finally, the chemical distance 𝑑*_chem_*(𝑄*_i_*, 𝑆*_i_*) corresponds to the previously introduced modified BLOSUM62 matrix (**Tab. S4**).

Overall, higher RUS*_i_* indicate residues that are chemically, spatially, and solvent-exposure distinct, and likely contribute to molecular specificity.

Residue similarity score (RSS) is derived from RUS, describing the degree of local conservation:

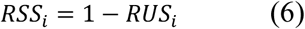

In pairwise one-to-one comparisons, uniqueness and similarity are strictly anti-correlated, but this is not necessarily true in one-to-many comparisons.

### One-to-many comparisons

In one-to-many comparisons, each query residue is evaluated against multiple subject proteins through successive one-to-one analyses. The minimum *RUS_i_* and *RSS_i_* scores observed across comparisons are then retained, providing the most stringent measure of conservation or divergence.

### Residue discriminability score

(RDS) calculations require at least two subjects, one (or more) “to engage” and one (or more) “to avoid”. RDS integrates RUS and RSS to identify residues that are distinct relative to some variants to be avoided (e.g. immunisation host proteins or non-target) and conserved relative to some other variants to engage (e.g. preclinical orthologues):

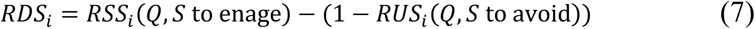

Negative RDS values, if any, are set to zero, as they represent residues with inverted discriminability.

### Combination of Residue Scores Across Surface Patches

Residue-level scores can be aggregated across predefined surface patches, such as known or putative epitopes, to provide region-level measures of divergence and conservation. Known epitopes characterised in this work were defined using crystallographic complex structures, where residues were included in the epitope if they contained at least one heavy atom within 4 Å of a binding-partner atom.

Surface selectivity, similarity, and discriminability scores were calculated as the mean of respectively RUS, RSS, and RDS across the patch under scrutiny. Aggregated patch-level scores enabled direct validation of the chemical, spatial, and solvent-exposure derived metrics against known binding interfaces and facilitated benchmarking of selectivity in relation to experimentally determined binding behaviour.

### Residue clustering to automatically identify putative epitopes

To automatically identify surface patches enriched in residues with high surface scores, which could be regarded as putative binding sites or epitopes more likely to yield the desired binding specificity, residue scores (RUS, RSS, or RDS) of the whole query protein were min-max normalised (0 to 1), and residues with values below 0.3 were excluded from further analysis. Each remaining residue was treated as a cluster seed, and neighbouring residues identified in the previous analysis that also exceeded the threshold were iteratively added to the same cluster. Clusters sharing at least one residue were merged to form a single contiguous region. Each cluster then corresponds to a putative epitope, and surface scores (as described above, without any min-max normalisation) are calculated to provide a representative value for the cluster. The top three clusters per analysis were selected for visualisation in some figures.

### Sequence- and Structure-Based Comparison for Benchmarking

To provide baseline comparisons, mean BLOSUM62 substitution scores^70^ were calculated across epitope residues for each query-subject pair. Epitope residues were extracted from the respective query PDB structure and the corresponding subject residues were assigned by sequence alignment (both pairwise alignment between the query and each subject, and a global multiple-sequence alignment comprising query and all subjects were used to calculate correlation coefficients in **Fig. 3j**). Alignments were carried out using the BLOSUM62 matrix itself for scoring, and default gap opening penalties of 10 and gap extension penalties of 0.5, from EMBL-EBI^71^. Query and subject amino acid sequences were extracted from PDB structures.

For pairwise sequence alignment, Needleman-Wunsch algorithm (global alignment) was used on extracted sequences. For multiple sequence alignment (MSA), sequences were aligned with the EMBL-EBI MUSCLE webserver (https://www.ebi.ac.uk/jdispatcher/msa/muscle)^71^.

For structural comparison, structural superimposition was achieved using TM-align algorithm^72,73^ on query and subject PDB structures before calculation of RMSD and TM-score on epitope residues.

### Correlations With Experimental Data

Binding affinities or functionality proxies, including IC₅₀, Kᵢ, and Kᴅ values, were sourced from the IUPHAR Guide to Pharmacology^40^ or peer-reviewed literature (see references in relevant sections of the text). Where affinity ranges were reported, mean values were used for the analysis, and error bars indicating the maximum and minimum of the reported range were plotted. Reported affinities were converted to logarithmic fold-changes relative to a reference protein (WT), producing a selectivity metric (i.e., 𝐿𝑜𝑔_10_(𝐾𝐷_variant_/ 𝐾𝐷_WT_) or *IC_50_*, or *K_i_*).

Pearson correlation coefficients (R) between SurfDiff scores and these logarithmic fold-changes were computed separately for each binder.

### Surface Mapping and Visualisation

Structural analyses and visualisations were conducted using PyMOL^74^, UCSF ChimeraX^75^, and in-house Python scripts built with Biopython, Matplotlib, and Seaborn. Residue-level scores were mapped onto protein surfaces using continuous colour gradients, with red indicating high uniqueness, blue representing high similarity, and white denoting low or intermediate values. Surface meshes (solvent accessible surface) were computed using the MSMS algorithm^76^ to provide high-resolution molecular surfaces suitable for score projection.

A custom-developed PyMOL plugin was used for streamlined visualisation and integration of residue scores into structural model meshes. This plugin enables direct loading, colouring, and exploration of annotated protein surfaces. Structural depictions of epitopes, binding interfaces, and divergence patterns were included in all visualisations to facilitate interpretation of residue-level and patch-level trends.

## Code availability

The SurfDiff source code is available at https://gitlab.developers.cam.ac.uk/ch/sormanni/surfdiff. A user-friendly webserver to run SurfDiff calculations is provided at www-cohsoftware.ch.cam.ac.uk/index.php/surfdiff. To access the webserver, users need to register a free account and log in.

## Author contributions

BJ and PS conceived the project. PS supervised the project. BJ, OW, and THN developed the code with the guidance of PS. BJ parsed the data and performed the calculations. BJ and PS critically analysed the results and wrote the first version of the paper. All authors contributed ideas, analysed data, and edited the paper. The contributions of the NIH authors are considered Works of the United States Government. The findings and conclusions presented in this paper are those of the authors and do not necessarily reflect the views of the NIH or the U.S. Department of Health and Human Services.

## Acknowledgements

P.S. is a Royal Society University Research Fellow (grant no. URF\R1\201461). We acknowledge funding from UK Research and Innovation (UKRI) Engineering and Physical Sciences Research Council (EPSRC grant no. EP/X024733/1, an ERC starting grant to P. S. underwritten by UKRI). Funding for THN was by the NIAID/NIH, NIH OxCam MD/PhD Scholars program, and the University of Wisconsin T32GM008692.

## Disclosure statement

The author(s) declare no potential conflict of interest.

## Declaration of usage of AI-assisted technologies

During the preparation of this work the authors used OpenAI GPT-5 to improve individual paragraph readability and shorten the text. GPT-4o was used to assist in optimising code. After using this tool/service, the author(s) reviewed and edited the content as needed, and take full responsibility for the content of the publication.

## Notes

### Competing Interest Statement

The authors have declared no competing interest.

