## Supplementary Figures and Tables for "SurfDiff: protein surface profiling for selective or broadly reactive epitope prioritisation in binder and immunogen design"

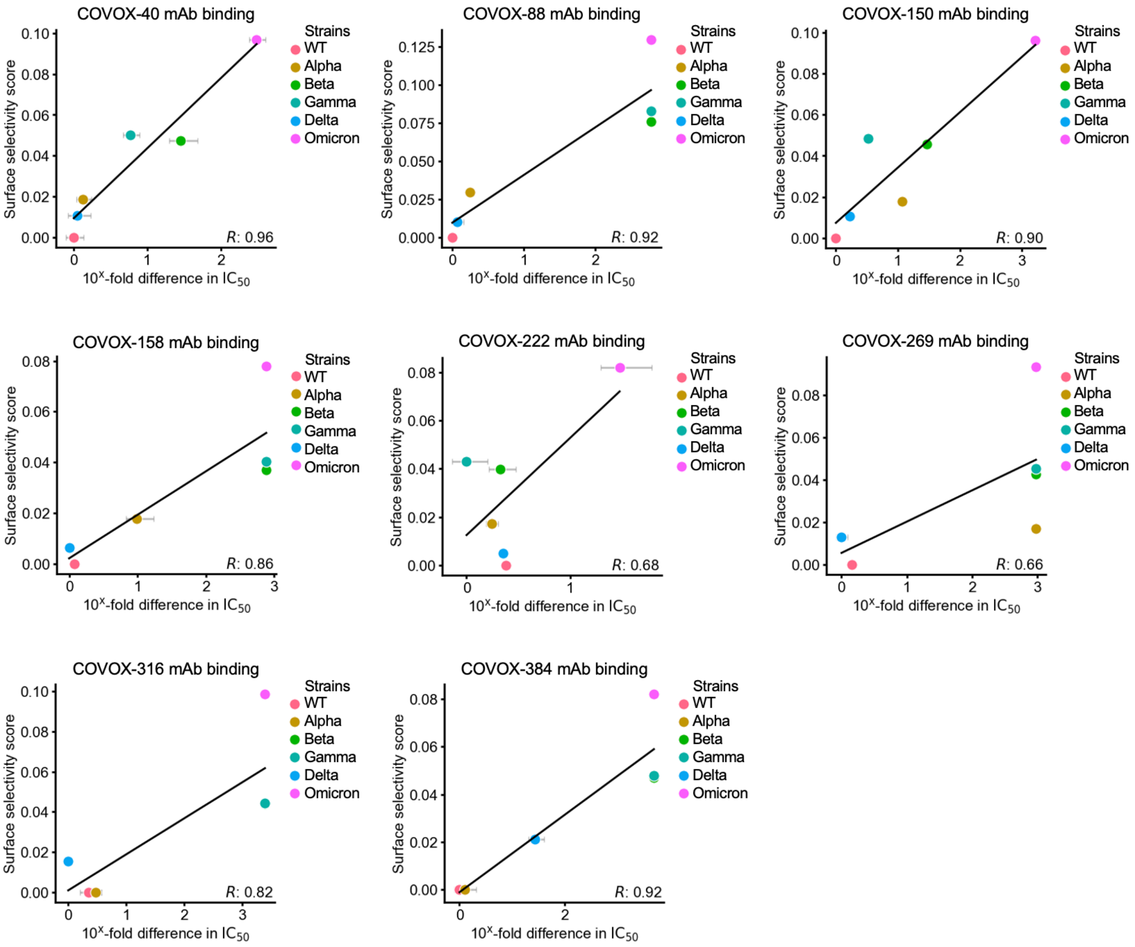


**Figure S1: Correlation between surface selectivity scores and antibody inhibitory potency differences across SARS-CoV-2 variants for individual COVOX antibodies.** Scatter plots show the relationship between SurfDiff surface selectivity scores and experimentally determined inhibitory potency differences (expressed as $\log_{10}(IC_{50}\_variant / IC_{50}\_WT)$) for each of the eight COVOX antibodies. Surface selectivity scores were computed over antibody epitopes defined from co-crystal structures using a 4 Å heavy atom distance cutoff. Each panel corresponds to a different antibody and presents the Pearson correlation coefficient (*R*) between surface selectivity and log-fold changes in neutralisation potency across SARS-CoV-2 variants. Variants that showed no neutralisation were assigned an $IC_{50}$ of 20 µg/mL, which is twice the reported detection limit.

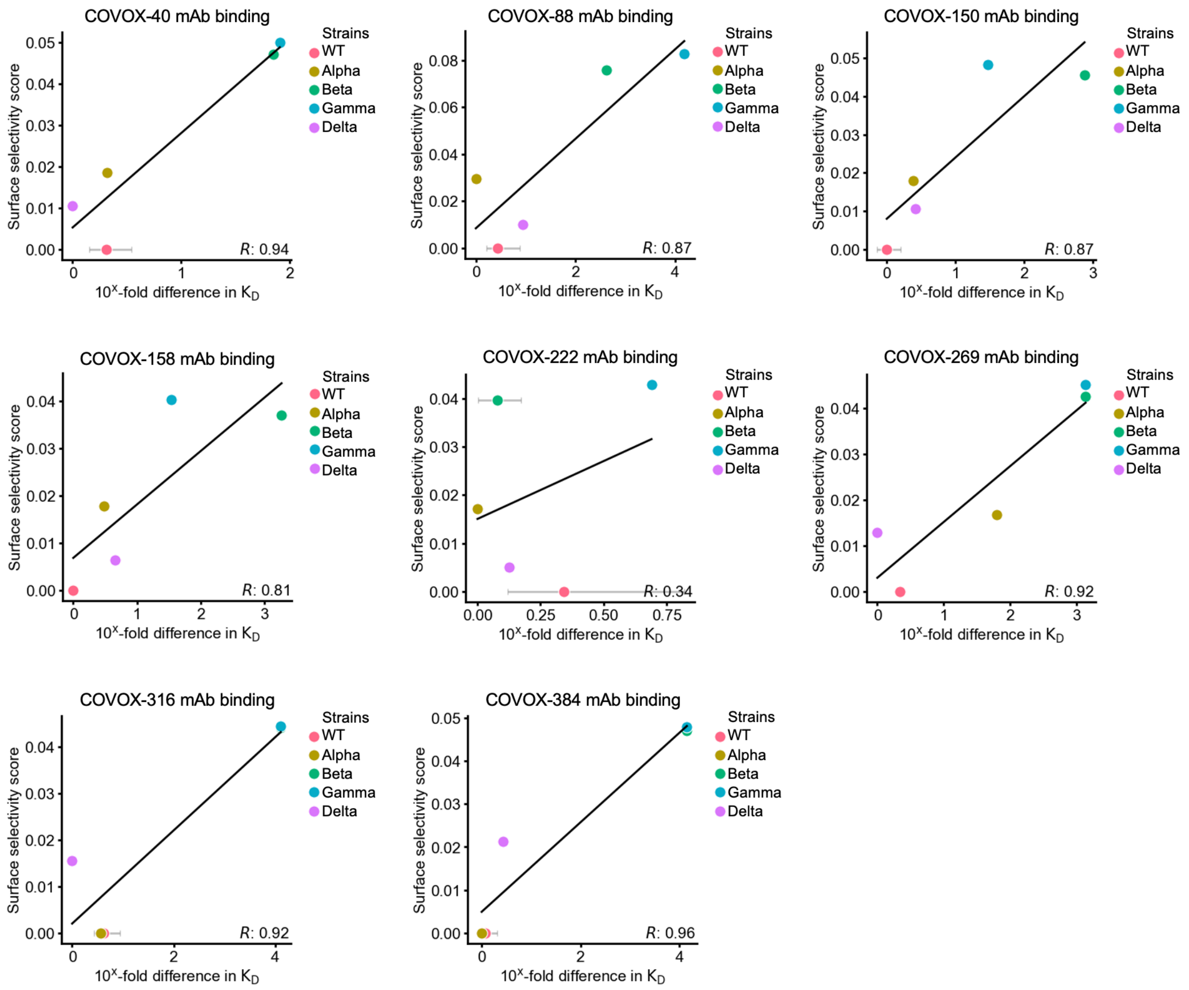


**Figure S2: Correlation between surface selectivity scores and antibody binding affinity differences across SARS-CoV-2 variants.** Scatter plots showing the relationship between SurfDiff surface selectivity scores and experimentally determined binding affinity differences ($\log_{10}(K_D\_variant\ /\ K_D\_WT)$) for each of the eight COVOX antibodies. Pearson correlation coefficients (*R*) are reported for each antibody. This panel is analogous to Supplementary Figure S1, but uses $K_D$ values instead of $IC_{50}$. Variants that showed no binding affinity were assigned a $K_D$ of 10 µM, which is one magnitude higher than the reported detection limit.

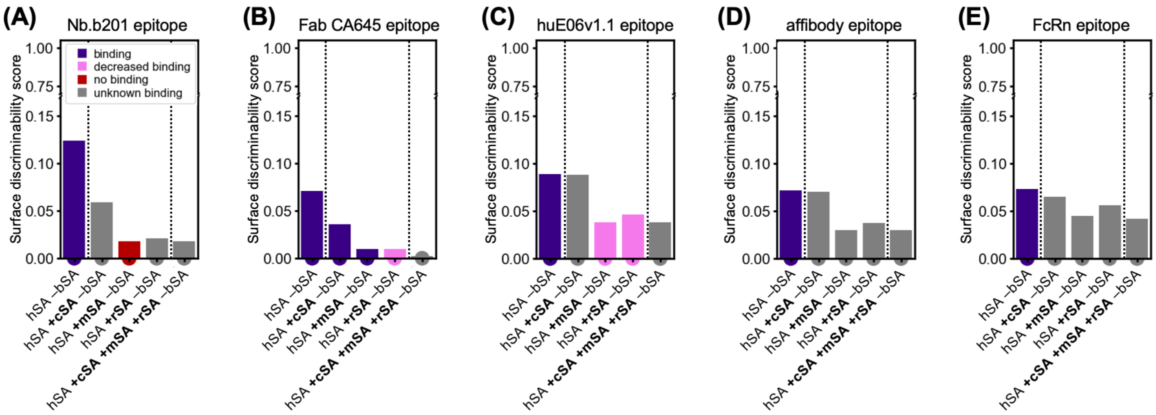


**Figure S3: Surface discriminability score analyses for multiple human serum albumin (hSA) binders.** Bar plots of surface discriminability scores for the epitopes of six experimentally characterised binders: nanobody Nb.b201, which was isolated with yeast display from a synthetic library (known to bind hSA and not mSA, while bSA was in the selection buffer so it's unlikely to bind that too). Fab CA645, shark VNAR huE06v1.1, an affibody (Affibody® ABD/GA), and the human neonatal Fc receptor (FcRn). Each panel displays the discriminability score at the respective epitope across hSA and its orthologues, enabling comparison of predicted cross-reactivity with reported experimental binding data.

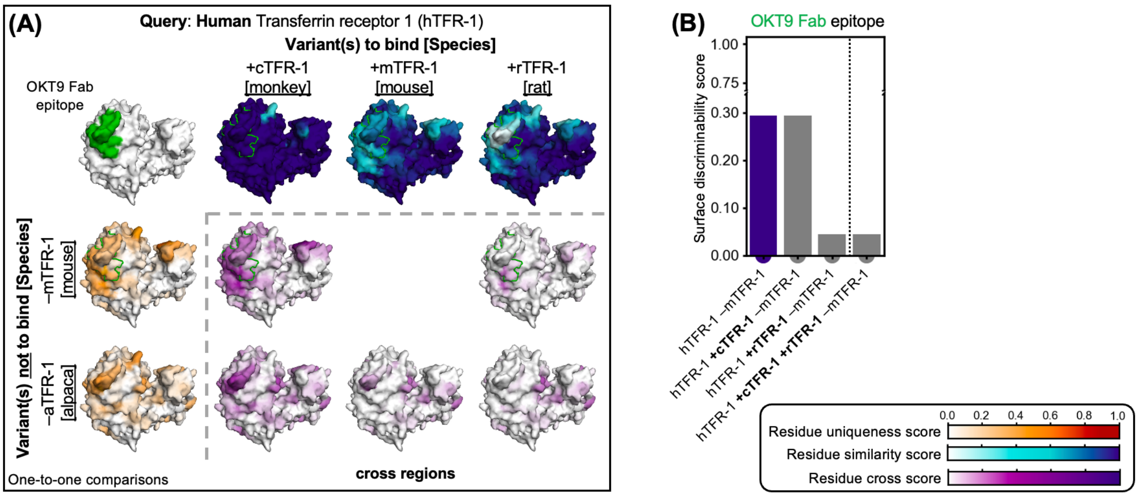


**Figure S4: Identification of potential cross-reactive epitopes across transferrin receptor 1 (TFR-1) orthologues using SurfDiff. (A)** Residue similarity and uniqueness scores mapped onto the surface of human TFR-1 (AlphaFold model, UniProt P02786) in comparison with orthologues from mouse, rat, alpaca, and cynomolgus monkey. Left: residue uniqueness scores relative to potential immunisation species (mouse, alpaca) highlight regions likely to elicit an immune response. Top: residue similarity scores relative to potential preclinical species (monkey, mouse, rat) highlight regions that may support cross-reactivity. Middle: surface discriminability scores integrate both metrics, identifying epitope regions that are distinct from immunisation hosts yet conserved across preclinical species. The OKT9 Fab epitope (AlphaFold3 model with docking-based orientation[1]) is shown in green.**(B)** Surface discriminability scores at the OKT9 epitope across variant(s) to bind. High scores for human are consistent with the known OKT9 binding profile, while low scores for mouse align with its lack of binding, reflecting development through mouse immunisation. A high score for monkey suggests potential cross-reactivity, although this has not been experimentally validated. In contrast, the low score for rat indicates limited predicted cross-reactivity, though experimental data are not available.

### SUPPLEMENTARY TABLES

| Cluster ID | Seed residue | Residues | Cluster size | Surface discriminability score [-] | Epitope SASA [Å$^2$] |
|---|---|---|---|---|---|
| 1 | 324 | [324, 325] | 2 | 0.37 | 236.0 |
| 2 | 528 | [528, 529] | 2 | 0.28 | 209.0 |
| 3 | 85 | [85, 88, 87] | 3 | 0.22 | 241.0 |
| 4 | 340 | [340, 344] | 2 | 0.21 | 97.0 |
| 5 | 141 | [141] | 1 | 0.19 | 92.0 |
| 6 | 585 | [585] | 1 | 0.18 | 25.0 |
| 7 | 96 | [96] | 1 | 0.15 | 56.0 |

**Table S1: Top-ranked epitope clusters on the surface of human serum albumin (hSA) with the highest surface discriminability scores for hSA +cSA +mSA +rSA –bSA –aSA.** Binding variants considered: hSA, cynomolgus monkey (cSA), mouse (mSA), and rat (rSA). Non-binding variants considered: alpaca serum albumin (aSA) and bovine serum albumin (bSA). Cluster information includes the cluster ID, the seed residue (UniProt residue numbering), all residues within the cluster (UniProt residue numbering), total number of residues, mean surface discriminability score, and solvent-accessible surface area (SASA). Query PDB: modelled with AF3 and signal peptide (residue 1-24) removed, UniProt ID: P02768).

| Cluster ID | Seed residue | Residues | Cluster size | Surface discriminability score [-] | Epitope SASA [Å$^2$] |
|---|---|---|---|---|---|
| 1 | 103 | [103] | 1 | 0.42 | 25.0 |
| 2 | 30 | [30, 31, 32] | 3 | 0.33 | 358.0 |
| 3 | 31 | [31, 30, 32, 135] | 4 | 0.28 | 404.0 |
| 4 | 68 | [68] | 1 | 0.27 | 35.0 |
| 5 | 32 | [32, 135, 31, 34, 30] | 5 | 0.26 | 468.0 |
| 6 | 133 | [32, 34, 36, 37, 38, 55, 133, 135] | 8 | 0.20 | 381.0 |
| 7 | 37 | [37, 36, 38, 53, 55] | 5 | 0.20 | 200.0 |
| 8 | 38 | [38, 36, 37, 53] | 4 | 0.20 | 174.0 |
| 9 | 53 | [53, 38, 55, 37] | 4 | 0.20 | 134.0 |

**Table S2: Top-ranked epitope clusters on the surface of human PD-1 (hPD-1) with the highest surface discriminability scores for hPD-1 +cPD-1 +rPD-1 –mPD-1.** Binding variants considered: hPD-1, cynomolgus monkey (cPD-1) and rat (rPD-1). Non-binding variants considered: mouse PD-1 (mPD-1). Cluster information includes the cluster ID, the seed residue (UniProt residue numbering), all residues within the cluster (UniProt residue numbering), total number of residues, mean surface discriminability score, and solvent-accessible surface area (SASA). Query PDB: 8gy5.

| Cluster ID | Seed residue | Residues | Cluster size | Surface discriminability score [-] | Epitope SASA [Å$^2$] |
|---|---|---|---|---|---|
| 1 | 127 | [127, 132] | 2 | 0.24 | 78.0 |
| 2 | 114 | [114, 116] | 2 | 0.14 | 221.0 |

**Table S3: Top-ranked epitope clusters on the surface of human PD-1 (hPD-1) with the highest surface discriminability scores.** Binding variants considered: hPD-1, cynomolgus monkey (cPD-1), mouse (mPD-1), and rat (rPD-1). Non-binding variants considered: alpaca PD-1 (aPD-1). Cluster information includes the cluster ID, the seed residue (UniProt residue numbering), all residues within the cluster (UniProt residue numbering), total number of residues, mean surface discriminability score, and solvent-accessible surface area (SASA). Query PDB: 8gy5.

| | | | | | | | | | | | | | | | | | | | | | |
|---|---|---|---|---|---|---|---|---|---|---|---|---|---|---|---|---|---|---|---|---|---|
| **A** | 0.00 | | | | | | | | | | | | | | | | | | | | |
| **R** | 0.67 | 0.00 | | | | | | | | | | | | | | | | | | | |
| **N** | 0.78 | 0.56 | 0.00 | | | | | | | | | | | | | | | | | | |
| **D** | 0.78 | 0.78 | 0.44 | 0.00 | | | | | | | | | | | | | | | | | |
| **C** | 0.56 | 0.83 | 0.78 | 0.78 | 0.00 | | | | | | | | | | | | | | | | |
| **Q** | 0.67 | 0.44 | 0.56 | 0.56 | 0.83 | 0.00 | | | | | | | | | | | | | | | |
| **E** | 0.67 | 0.56 | 0.56 | 0.33 | 0.89 | 0.33 | 0.00 | | | | | | | | | | | | | | |
| **G** | 0.56 | 0.78 | 0.56 | 0.67 | 0.78 | 0.78 | 0.78 | 0.00 | | | | | | | | | | | | | |
| **H** | 0.78 | 0.56 | 0.50 | 0.61 | 0.78 | 0.56 | 0.56 | 0.72 | 0.00 | | | | | | | | | | | | |
| **I** | 0.67 | 0.94 | 0.89 | 0.89 | 0.67 | 0.94 | 0.94 | 1.00 | 0.89 | 0.00 | | | | | | | | | | | |
| **L** | 0.67 | 0.78 | 0.89 | 1.00 | 0.67 | 0.78 | 0.94 | 1.00 | 0.89 | 0.28 | 0.00 | | | | | | | | | | |
| **K** | 0.67 | 0.33 | 0.56 | 0.67 | 0.83 | 0.44 | 0.44 | 0.78 | 0.67 | 0.94 | 0.78 | 0.00 | | | | | | | | | |
| **M** | 0.67 | 0.67 | 0.78 | 0.89 | 0.67 | 0.56 | 0.78 | 0.89 | 0.72 | 0.44 | 0.33 | 0.67 | 0.00 | | | | | | | | |
| **F** | 0.78 | 0.89 | 0.83 | 0.83 | 0.72 | 0.89 | 0.89 | 0.83 | 0.61 | 0.56 | 0.56 | 0.89 | 0.56 | 0.00 | | | | | | | |
| **P** | 0.67 | 0.72 | 0.72 | 0.67 | 0.78 | 0.67 | 0.67 | 0.72 | 0.72 | 0.89 | 0.89 | 0.67 | 0.72 | 0.89 | 0.00 | | | | | | |
| **S** | 0.44 | 0.67 | 0.44 | 0.56 | 0.67 | 0.56 | 0.56 | 0.56 | 0.67 | 0.83 | 0.83 | 0.56 | 0.67 | 0.78 | 0.67 | 0.00 | | | | | |
| **T** | 0.56 | 0.67 | 0.56 | 0.67 | 0.67 | 0.67 | 0.67 | 0.78 | 0.72 | 0.67 | 0.67 | 0.67 | 0.67 | 0.78 | 0.67 | 0.44 | 0.00 | | | | |
| **W** | 0.83 | 0.78 | 0.83 | 0.83 | 0.67 | 0.72 | 0.78 | 0.72 | 0.67 | 0.83 | 0.72 | 0.78 | 0.61 | 0.50 | 0.83 | 0.83 | 0.72 | 0.00 | | | |
| **Y** | 0.78 | 0.72 | 0.72 | 0.83 | 0.72 | 0.67 | 0.72 | 0.83 | 0.39 | 0.67 | 0.67 | 0.72 | 0.67 | 0.28 | 0.78 | 0.78 | 0.72 | 0.44 | 0.00 | | |
| **V** | 0.56 | 0.94 | 0.89 | 0.89 | 0.67 | 0.78 | 0.78 | 0.89 | 0.89 | 0.11 | 0.44 | 0.78 | 0.44 | 0.67 | 0.78 | 0.83 | 0.56 | 0.83 | 0.67 | 0.00 | |
| **-** | 1.00 | 1.00 | 1.00 | 1.00 | 1.00 | 1.00 | 1.00 | 1.00 | 1.00 | 1.00 | 1.00 | 1.00 | 1.00 | 1.00 | 1.00 | 1.00 | 1.00 | 1.00 | 1.00 | 1.00 | |
| **Aa** | **A** | **R** | **N** | **D** | **C** | **Q** | **W** | **G** | **H** | **I** | **L** | **K** | **M** | **F** | **P** | **S** | **T** | **W** | **Y** | **V** | **-** |

**Table S4. Residue Chemical Distance matrix used in SurfDiff calculations.** This matrix was derived from a modified BLOSUM62 substitution matrix reformulated to represent physicochemical distances rather than evolutionary background frequencies. Diagonal values (self-matches) were set to zero to isolate identical substitutions. The resulting scores were min–max normalised to span the range from 0 to 1, providing a scale interpretable as physicochemical dissimilarity for each residue (Aa). Residues classified as different (–) were assigned the maximum score of 1.00.

| | **PDB-ID / AF3 (UniProt-ID)** | **Binding complex with** |
|---|---|---|
| **SARS-CoV-2 (RBD) strains** | | |
| WT [Victoria] | 7neh / 7z0x / 7nd3 / 7bel / 7bei / 7qny / 7nx6 / 7beh / 7bep | COVOX-40 (7nd3), -88 (7bel), -150 (7bei), -158 (7qny), -222 (7nx6), -269 (7neh), -316 (7beh), -384 (7peb) |
| Alpha [B.1.1.7] | 7ek0 | |
| Beta [B.1.351] | 7ps7 | |
| Gamma [P.1] | 7nxc | |
| Delta [B.1.617.2] | 7x7d | |
| Omicron [B.1.1.529] | 7qnw | |
| **Serotonin receptor subtypes** | | |
| 5-HT1A | 7e2y / 7e2z | serotonin (7e2y) |
| 5-HT1B | 6g79 | |
| 5-HT1D | 7e32 | |
| 5-HT1E | 7e33 | |
| 5-HT1F | 7exd | |
| 5-HT2A | 7ran / 8uwl | lisuride (8uwl) |
| 5-HT2B | 7srr | |
| 5-HT2C | 8dpf | |
| 5-HT4 | 7xt8 | |

| | | |
|---|---|---|
| 5-HT5A | 7um7 | |
| 5-HT6 | 8jlz | |
| 5-HT7 | 7xtc | 5-carboxamido-tryptamine |
| **Dopamine receptor subtypes** | | |
| D1 | 7jvp | |
| D2 | 7jvr | |
| D3 | 7cmv | |
| D4 | 8iru | |
| D5 | 8irv | |
| **Adrenergic receptor subtypes** | | |
| Aa1A | 8thk | |
| Aa1B | 7b6w | |
| Aa1D | AF3 (P25100) | |
| Aa2A | 7ej8 | |
| Aa2B | 6k41 | |
| Aa2C | 6kuw | |
| Ab1 | 7bu6 | |
| Ab2 | 4lde | |
| Ab3 | 9ijd | |
| **Vasopressin / Oxytocin receptor subtypes** | | |
| V1A | AF3 (P37288) | |
| V1B | AF3 (P47901) | |
| V2 | 7dw9 | vasopressin |
| OT | 7qvm | oxytocin |
| **Somatostatin receptor subtypes** | | |
| SSTR1 | 8xio | |
| SSTR2 | 7t10 | somatostatin-14 |
| SSTR3 | 8xir | |
| SSTR4 | 7xms | |
| SSTR5 | 8x8n | |
| **Dengue virus (DIII envelope protein) strains** | | |
| DENV-1 [FGA89] | 3uzq / 4l5f | mAb 4E11 (3uzq) |
| DENV-2 [1409] | 8y3k | |
| DENV-3 [PaH881] | 8jn4 | |
| DENV-4 [63632] | 3uyp | |
| **HIV (gp120 envelope protein) strains** | | |
| HIV-1 [Q23.17] | 4ydi | |
| HIV-1 [KER_2018_11] | 4lss | |
| HIV-1 [Q842.d12] | 4xmp | |
| HIV-1 [93TH057] | 5te6 / 4xvt | mAb N6 (5te6) |
| HIV-1 [YU2] | 3tgg | |
| HIV-1 [ZM176.66] | 4lst | |
| HIV-1 [C1086] | 4lsv | |
| HIV-1 [DU172.17] | 5te7 | |
| HIV-1 [X2088] | 5te4 | |
| HIV-2 | 5cay | |
| **AngiotensinII receptor subtypes** | | |
| ATR1 | 6os2 | |
| ATR2 | 6jod / 5xjm | AngII (6jod) / 4A03 Fab (6jod) |
| **Transforming growth factor-beta cytokine isoforms** | | |

| | | |
|---|---|---|
| TGF-β1 | 4kv5 | fresolimumab |
| TGF-β2 | 4kxz | fresolimumab |
| TGF-β3 | 3eo1 | fresolimumab |
| **Serum Albumin species orthologs** | | |
| Human [homo sapiens] | AF3 (P02768) / 8oi2 / 5vnw / 5fuo / 1tf0 / 4n0f | NbAlb1 (8oi2), Nb.b201 (5vnw), Fab CA645 (5fuo), affibody (1tf0), FcRn (4n0f) |
| Monkey [cynomolgus monkey] | AF3 (A2V9Z4) | |
| Mouse [mus musculus] | AF3 (P07724) | |
| Rat [rattus norvegicus] | AF3 (P02770) | |
| Bovine [bos taurus] | AF3 (P02769) | |
| Alpaca [lama pacos] | AF3 (A0A6I9IHM8) | |
| Shark | 4hgm | huE06v1.1 |
| **PD-1 species orthologs** | | |
| Human [homo sapiens] | 8gy5 | cemiplimab |
| Monkey [cynomolgus monkey] | AF3 (Q4R367) | |
| Mouse [mus musculus] | 3bik | |
| Rat [rattus norvegicus] | AF3 (A6JR44) | |
| Alpaca [lama pacos] | AF3 (A0A6J0ASH2) | |
| **Transferrin receptor 1 species orthologs** | | |
| Human [homo sapiens] | AF3 (P02786) | OKT9 Fab (as modelled in[1]) |
| Monkey [cynomolgus monkey] | AF3 (A0A2K5X958) | |
| Mouse [mus musculus] | AF3 (Q62351) | |
| Rat [rattus norvegicus] | AF3 (Q99376) | |
| Alpaca [lama pacos] | AF3 (A0A6J0ACT4) | |
| **Negative controls** | | |
| GLP-1R | 6x18 | |
| GFP | 5b61 | |
| RNase A | 7rsa | |
| HSA | 1ao6 | |

**Table S5: Overview of protein targets, structural models, and binding complexes used for analysis and benchmarking.** The table lists the protein systems investigated in this study, including negative control proteins. For each protein or variant, the corresponding molecular structures are given as either experimentally determined structures (PDB IDs) or AlphaFold-predicted structural models (AF3, with UniProt accession numbers in parentheses). Where available, used structures of the protein in complex with a known binder – such as an antibody, ligand, peptide, or Fab fragment – are listed, along with the identity of the interacting molecule. These complexes were used for annotation of binding epitopes, which were exploited for validating surface selectivity, similarity, and discriminability scores. Species orthologs and viral strain variants are indicated where relevant. Negative controls represent structurally unrelated proteins without known binding relevance to the systems under investigation. The used query structure/model is underlined.
